# An approximate likelihood method reveals ancient gene flow between human, chimpanzee and gorilla

**DOI:** 10.1101/2023.07.06.547897

**Authors:** Nicolas Galtier

## Abstract

Gene flow and incomplete lineage sorting are two distinct sources of phylogenetic conflict, *i*.*e*., gene trees that differ in topology from each other and from the species tree. Distinguishing between the two processes is a key objective of current evolutionary genomics. This is most often pursued via the so-called ABBA-BABA type of method, which relies on a prediction of symmetry of gene tree discordance made by the incomplete lineage sorting hypothesis. Gene flow, however, need not be asymmetric, and when it is not, ABBA-BABA approaches do not properly measure the prevalence of gene flow. I introduce Aphid, an approximate maximum-likelihood method aimed at quantifying the sources of phylogenetic conflict via topology and branch length analysis of three-species gene trees. Aphid draws information from the fact that gene trees affected by gene flow tend to have shorter branches, and gene trees affected by incomplete lineage sorting longer branches, than the average gene tree. Accounting for the among-loci variance in mutation rate and gene flow time, Aphid returns estimates of the speciation times and ancestral effective population size, and a posterior assessment of the contribution of gene flow and incomplete lineage sorting to the conflict. Simulations suggest that Aphid is reasonably robust to a wide range of conditions. Analysis of coding and non-coding data in primates illustrates the potential of the approach and reveals that a substantial fraction of the human/chimpanzee/gorilla phylogenetic conflict is due to ancient gene flow. Aphid also predicts older speciation times and a smaller estimated effective population size in this group, compared to existing analyses assuming no gene flow.

## 1 Introduction

Phylogenetic conflicts occur when distinct genes support distinct trees. Conflicts often result from errors, when a reconstructed tree does not reflect the true history of the considered locus. Hidden paralogy, alignment problems and substitution model mis-specification are, among others, well-documented sources of discrepancy between gene trees and species trees (Scornavacca et al., 2020). These problems are presumably mitigated when closely related species are analyzed. In this case, it is likely that a majority of observed conflicts reflect genuine differences among genealogies. Two biological processes are potentially involved: gene flow (GF) and incomplete lineage sorting (ILS).

GF, or introgression, happens when a piece of DNA is transferred from one species to another. If the immigrant allele spreads and reaches fixation, then the previously accumulated divergence between the donor and recipient species is erased, and the two species get to be genealogically related at this locus, regardless of the speciation history. Loci experiencing GF are therefore likely to have genealogies that differ from each other, and from loci unaffected by GF. GF has been documented in a wide range of taxa (Green et al., 2010; Abby et al., 2012; Fontaine et al., 2015; Meyer et al., 2017; Ropars et al., 2018; Edelman et al., 2019; Glémin et al., 2019; Zhang et al., 2021; Suvorov et al., 2022b), and in eukaryotes typically happens via hybridization between closely related species, when reproductive isolation is not yet complete.

ILS is an independent source of phylogenetic conflict that happens when within-species polymorphism lasts longer than the time between two successive events of speciation. Consider for instance that, at one particular locus, three alleles, *X*_1_, *X*_2_ and *X*_3_, segregate in an ancestral species. Imagine that the ancestral species quickly splits twice, and that all three alleles are transmitted to the newly formed species *A, B* and *C*. Now *A, B* and *C* start diverging, and each of them randomly keeps one of *X*_1_, *X*_2_ or *X*_3_ by drift. The phylogeny at this locus will reflect the pre-existing genealogical relationships between *X*_1_, *X*_2_ and *X*_3_ in the ancestral species, not the sequence of speciation events giving rise to *A, B* and *C*. The expected prevalence of ILS depends on the time between speciation events and on the effective population size (Hudson, 1983). ILS has been documented in a number of taxa, including great apes (Mallet et al., 2016; Dutheil et al., 2009; Hobolth et al., 2011; Guerzoni and McLysaght, 2016; Mendes and Hahn, 2016; Meleshko et al., 2021; Rivas-González et al., 2023a).

GF and ILS are two different processes, each responsible for a fraction of the genuine variation between gene trees. A natural question to ask is the relative contribution of ILS and GF to the observed conflict, which raises the issue of how to distinguish between the two processes. The literature on this subject is rich and diverse (Jiao et al., 2021; Hibbins and Hahn, 2022). To model phylogenies with ILS is relatively straightforward. The Multi-Species Coalescent model assumes that loci have independent coalescence histories within the framework of a shared species tree, speciation times, and effective population size (Rannala and Yang, 2003). Inference methods based on this model have been developed and applied to numerous data sets (Rannala and Yang, 2017), including a modified version also accounting for the auto-correlation of genealogies along chromosomal segments (Dutheil et al., 2009). Adding GF to the picture is a more challenging task. Recent developments have been made in this direction, both theoretical and in terms of inference tools (Long and Kubatko, 2018; Wen and Nakhleh, 2018; Glémin et al., 2019; Flouri et al., 2020; Rogers, 2019, 2022). Some of these methods in principle fulfil the objective of jointly estimating the prevalence of ILS and GF. These methods, however, are computationally demanding, and/or require specific hypotheses about which lineages have exchanged genes. They have not been applied to a large number of data sets so far.

Rather, the empirical literature is dominated by a hypothesis-testing approach, in which the existence of GF is demonstrated via rejection of an ILS-only scenario. This is based on a prediction made by the ILS model, which is that, if ((*A, B*), *C*) is the true species tree, the alternative ((*A, C*), *B*) and ((*B, C*), *A*) trees should be observed in roughly equal frequencies. GF, in contrast, can be asymmetric, *i*.*e*., there could be more gene flow between *A* and *C* than between *B* and *C*, for instance, leading to a larger number of ((*A, C*), *B*) than ((*B, C*), *A*) gene trees. A significant imbalance between the discordant topologies therefore demonstrates the existence of GF. The popular ABBA-BABA test (Green et al., 2010; Durand et al., 2011) and its variations (Reich et al., 2010; Meyer et al., 2012; Pease and Hahn, 2015; Blischak et al., 2018) implement this idea. These methods are efficient and have routinely been applied to a large number of data sets.

It should be noted that, although perfectly sound for detecting the existence of GF, ABBA-BABA-like approaches are not suitable for quantifying its prevalence. This is because GF does not need to be asymmetric, but only its asymmetric component is detected by ABBA-BABA. If GF between C and A (on one hand) and between C and B (on the other hand) occurred at a similar rate, then ABBA-BABA approaches would simply have no power (Vanderpool et al., 2020). Yet, even in the case of symmetric gene flow, the ILS and GF hypothesis can in principle be distinguished since they make distinctive predictions regarding coalescence times. Indeed, under GF the time to the most recent common ancestor of the interacting lineages is reduced compared to non-GF scenarios, whereas ILS, in contrast, entails relatively old coalescences. This means that besides topology counts, one can potentially gain information on the prevalence of GF and ILS by examining gene tree branch lengths. Although present in the literature (Holder et al., 2001; Joly et al., 2009; Smith and Kronforst, 2013; Meyer et al., 2017; Suvorov et al., 2022a), this idea has rarely been used as a basis for of method distinguishing GF from ILS. A notable exception is the recently developed QuIBL method (Edelman et al., 2019), which focuses on the distribution of the length of the internal branch in three-species rooted gene trees. The ILS hypothesis predicts an exponential distribution for this variable for trees discordant with the species tree, whereas in the case of GF, gene trees departing the species tree topology should be associated with an increased mean and variance of the internal branch length.

Of note, not only the internal but also terminal branch lengths differ in expectation under an ILS *vs*. a GF scenario (Fig.1). Here I introduce a new method, Aphid, aiming at capturing this information. Aphid basically asks whether discordant gene trees tend to be taller, indicating ILS, or shorter, indicating GF, than the majority gene trees. Modeling a set of three-species rooted gene trees as a mixture of no-event, ILS and GF scenarios, Aphid considers both terminal and internal branch lengths and accounts for differences in mutation rate among loci, while making a number of simplifications that render it particularly efficient. Analysis of simulated and real data sets shows that, unlike ABBA-BABA tests, Aphid can reveal the existence of symmetric GF during species divergence. In particular, Aphid predicts that a substantial portion of the phylogenetic conflict in great apes results from ancient GF between the human, chimpanzee and gorilla lineages.

## 2 The method

### 2.1 Mixture of genealogies

Consider a triplet of species ((*A, B*), *C*) such that the (*A, B*)|*C* split occurred *t*_2_ generations ago and the *A*|*B* split occurred *t*_1_ generations ago, and assume a constant population size of *N*_*e*_ diploid individuals across the whole tree. Suppose that we sample exactly one haplome per species and reconstruct the genealogical history (gene tree) of a number of genomic segments (loci). This set of gene trees is modeled as coming from a limited mixture of characteristic trees here called scenarios, which have fixed branch lengths set to their expected values given the scenario. These fall into three categories: no-event, ILS, and GF. The no-event category corresponds to the scenario in which neither GF nor ILS has been involved (Fig.1, scenario *S*_1_). Under this scenario the tree topology must be ((*A, B*), *C*), and coalescence times are assumed to equal *t*_1_ + 2*N*_*e*_ and *t*_2_ + 2*N*_*e*_ for the *A*|*B* and (*A, B*)|*C* split, respectively, 2*N*_*e*_ being the average coalescence time between two lineages in a panmictic population. The ILS category corresponds to three scenarios in which the coalescence between the *A* and *B* lineages is older than *t*_2_. In this case the three possible topologies can be generated, hence scenario *S*_2_, *S*_3_ and *S*_4_ in Fig.1. The coalescence times are modeled to be fixed at *t*_2_ + 2*N*_*e*_*/*3 and *t*_2_ + 2*N*_*e*_*/*3 + 2*N*_*e*_, respectively, 2*N*_*e*_*/*3 being the average time of the first coalescence among three lineages in a panmictic population. Finally, the GF category corresponds to scenarios in which an event of gene flow has occurred. Fig.1 displays five such scenarios, depending on which lineage is the donor and which the recipient. Scenario *S*_5_ involves an event of gene flow between the *A* and *B* lineages. This does not affect the topology, but leads to a younger *A*|*B* coalescence time compared to scenario 1. Scenario *S*_6_ and *S*_7_ involve a transfer from *A* to *C* and *C* to *A*, respectively, which both result in an ((*A, C*), *B*) topology while also modifying the expected coalescence times and branch lengths. Scenario *S*_8_ and *S*_9_ similarly model gene flow between *B* and *C*.

Branch lengths in GF scenarios depend on the *t*_*g*_ parameter, which represents the coalescence time of two lineages brought together by gene flow — *i*.*e*., on average, GF time plus 2*N*_*e*_. The five GF scenarios shown in Fig.1 are such that the same GF coalescence time is shared by all loci experiencing GF. Our method actually relaxes this assumption by assuming a discrete distribution, instead of fixed values, of GF coalescence time. The user is asked to choose a finite number, *n*_*c*_, of times, expressed as fractions of *t*_1_. The oldest of these GF coalescence times is assigned probability *p*_*a*_ (for “ancient”), while each of the others has probability (1 − *p*_*a*_)*/*(*n*_*c*_ − 1). In this study, the default setting was *n*_*c*_ = 2 and it was assumed that the coalescence of lineages brought together by GF events could occur at time *t*_1_ or *t*_1_*/*2. Ten GF scenarios were therefore considered, as explicitly described in Supplementary Fig. 1 and Supplementary Table 1.

**Figure 1.**
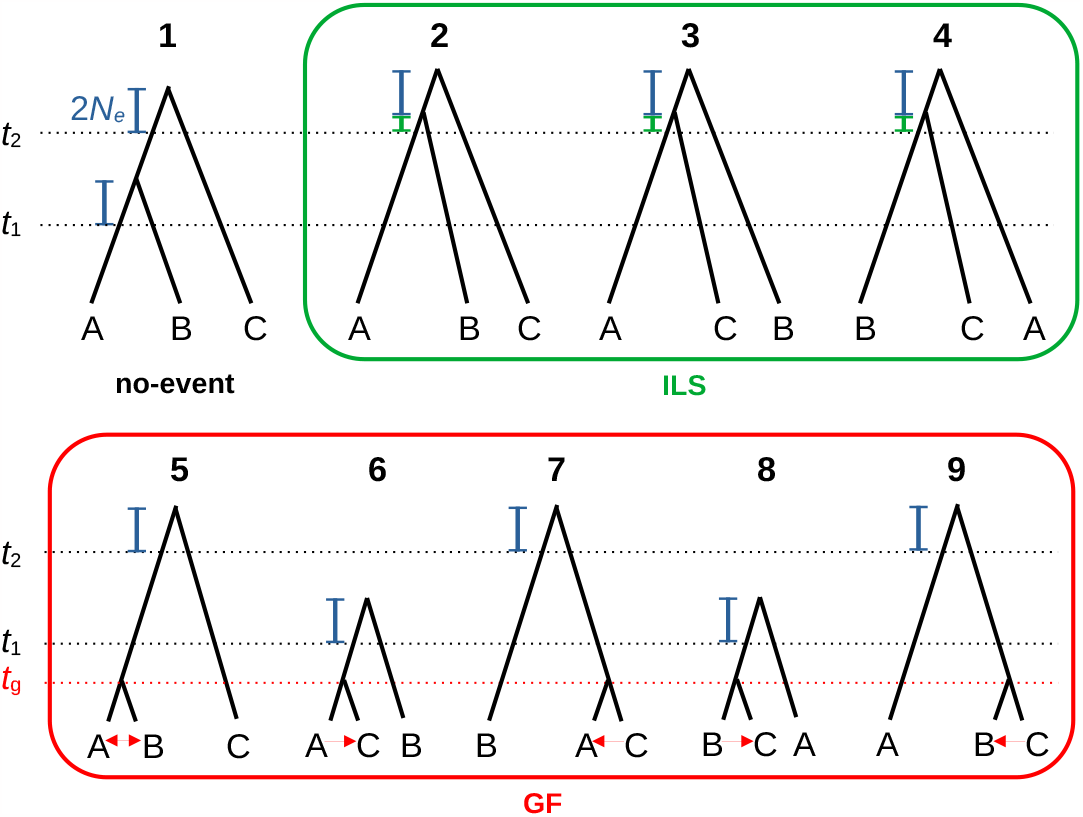
Typical scenarios involving incomplete lineage sorting (green), gene flow (red) or no event. Branch lengths depend on speciation times *t*_1_ and *t*_2_ and effective population size *N*_*e*_. The vertical blue and green bars are of length 2*N*_*e*_ and 2*N*_*e*_*/*3, respectively. Observed gene trees are modelled as a mixture of such scenarios. See Supplementary Figure S1 and Supplementary Table 1 for a detailed version.

Our model assigns prior probabilities to these scenarios. GF between *A* and *B, A* and *C*, and *B* and *C* are assumed to occur with probability *p*_*AB*_, *p*_*AC*_ and *p*_*BC*_, respectively. GF is here assumed to be bidirectional, and to have the same probability to occur from *A* to *C* as from *C* to *A*, and from *B* to *C* as from *C* to *B* — events of GF from *A* to *B* and from *B* to *A* lead to the very same pattern and are indistinguishable anyway. The *p*_*AB*_, *p*_*AC*_ and *p*_*BC*_ parameters, combined with *p*_*a*_, define the prior probabilities of the ten GF scenarios, as detailed in Supplementary Table 1. In the absence of gene flow, ILS will occur if the *A* and *B* lineages have not yet coalesced at time *t*_2_, which happens with probability 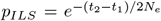 (Hudson, 1983). *Each of the three ILS scenarios therefore has probability (1* − *p*_*AB*_ − *p*_*AC*_ − *p*_*BC*_)*p*_*ILS*_*/*3. The no-event scenario, finally, has probability (1 − *p*_*AB*_ − *p*_*AC*_ − *p*_*BC*_)(1 − *p*_*ILS*_). This parametrization entails a constraint on the sum of GF probabilities, *p*_*AB*_ + *p*_*AC*_ + *p*_*BC*_, which must be lower than one.

### 2.2 Mutation rate

The above description expresses time in generation units, whereas what we can measure from the data is sequence divergence, which for a neutral locus depends on coalescence time and mutation rate. Calling *μ* the locus mutation rate, the *t*_1_, *t*_2_ and *N*_*e*_ parameters are re-scaled as:

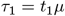

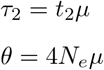

Crucially, the method accounts for the variation in mutation rate among loci. This is required if one aims at interpreting differences in branch length among gene trees in terms of GF or ILS. A locus-specific factor, *α*_*i*_, is introduced, corresponding to the relative mutation rate of the *i*^*th*^ locus and multiplying the predicted branch lengths for this locus. Information on *α*_*i*_’s will be obtained by using additional species besides *A, B* and *C* (see below). Supplementary Table 1 expresses branch lengths as functions of the re-scaled parameters under the 14 scenarios.

### 2.3 Parameter estimation

The model includes seven parameters controlling the distribution of gene tree topology and branch lengths (speciation times *τ*_1_ and *τ*_2_, population mutation rate *θ*, gene flow probabilities *p*_*AB*_, *p*_*AC*_, *p*_*BC*_, *p*_*a*_) and one relative mutation rate per locus, *α*_*i*_.

The whole set of parameters is not identifiable from three-species gene trees only since the predicted branch lengths are expressed as products of *α*_*i*_ and *τ*_1_, *τ*_2_, or *θ* (Supplementary Table 1). An external source of information on the *α*_*i*_’s is therefore needed. Aphid requires that every input gene tree includes at least one species from a pre-specified outgroup clade, in addition to *A, B* and *C*. The relative mutation rate at locus *i, α*_*i*_, is estimated as 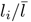, where *l*_*i*_ is the mean distance from root to tip in the *i*^th^ gene tree, and 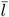 is the average of *l*_*i*_’s across gene trees. This relies on the assumptions that the mutation rate is constant in time and among lineages, *i*.*e*., that sequences evolve in a clock-wise manner.

The other seven parameters are estimated in the maximum likelihood framework. The probability of the *i*^th^ gene tree given the parameters is written as:

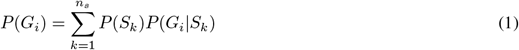

where *G*_*i*_ is the *i*^th^ gene tree, *S*_*k*_ the *k*^th^ scenario, and *n*_*s*_ the total number of scenarios (here *n*_*s*_ = 14). The prior probabilities of the various scenarios, *P* (*S*_*k*_), are given in Supplementary Table 1 and detailed in the last paragraph of section 2.1 above. The probability of a gene tree given a scenario, *P* (*G*_*i*_|*S*_*k*_) equals zero if the topology of *G*_*i*_ does not match *S*_*k*_, and is written as:

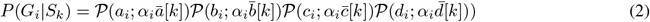

if the topology of *G*_*i*_ matches *S*_*k*_. Here, *a*_*i*_, *b*_*i*_, *c*_*i*_ and *d*_*i*_ are the terminal branch lengths of the *A, B* and *C* lineages and internal branch in *G*_*i*_, respectively, 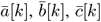 and 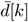] are the expected branch lengths under scenario *S*_*k*_ (Supplementary Table 1), and *𝒫* (*j*; *λ*) = *λ*^*j*^*e*^−*λ*^*/j*! is the probability mass function of the Poisson distribution. Note that branch lengths are here defined as the total, not per site, number of sequence changes having occurred between two nodes. The method can easily account for unresolved (*i*.*e*., star-like) gene trees: the probability of an unresolved gene tree given a scenario is given by equation (2) irrespective of scenario topology. Finally, the likelihood of the whole data set is obtained by multiplying the likelihood across gene trees, here assumed to be independent. The likelihood was maximized using an adapted version of the Newton-Raphson algorithm, which relies on differentiation of the likelihood function with respect to the parameters.

### 2.4 Gene tree annotation, GF/ILS contribution, imbalance

Having estimated the parameters, one can calculate the posterior probability of each of the scenarios for any particular gene tree using an empirical Bayesian approach:

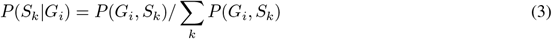

The posterior probability for a gene tree to be affected by ILS can be calculated as:

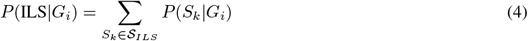

and similarly for GF:

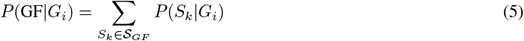

where *𝒮*_*LS*_ and *𝒮*_*GF*_ are the sets of ILS and GF scenarios, respectively.

The estimated contributions of ILS and GF to the phylogenetic conflict in the whole data set are expressed as:

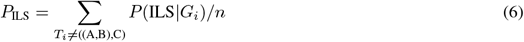

and

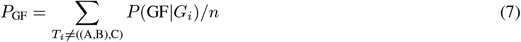

where *T*_*i*_ is the topology of gene tree *G*_*i*_ and *n* is the number of analyzed gene trees. Equation 6 and 7 sum posterior probabilities across all gene trees having a discordant topology, either ((*A, C*), *B*) or ((*B, C*), *A*). Separating these two terms allows one to define the discordant topology imbalance associated with ILS as

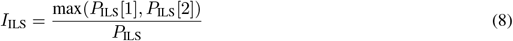

where

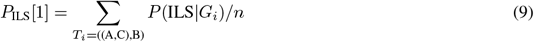

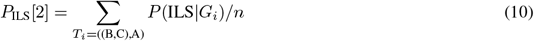

and similarly for GF:

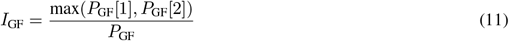

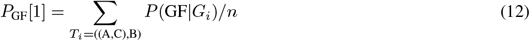

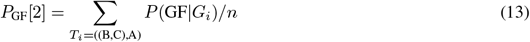

*I*_ILS_ and *I*_GF_ vary between 0.5 (no imbalance) and 1 (maximal imbalance). *I*_GF_ measures the difference in the amount of GF between A and C (on one hand) and between B and C (on the other hand), in the spirit of ABBA-BABA. *I*_ILS_ is expected to be close to 0.5, ILS being a symmetrical process. A value of *I*_ILS_ much larger than 0.5 might indicate that Aphid failed in capturing the true population history, and/or reflect the existence of GF from so-called “ghost” lineages (see discussion).

### 2.5 Confidence interval, hypothesis testing

95% confidence intervals around maximum likelihood parameter estimates were defined as intervals including all values of the considered parameter such that the log-likelihood dropped off by less than 1.92 units. These were calculated by re-optimising the likelihood over all the parameters other than the one being investigated for each test value of the parameter under consideration. Likelihood ratio tests were conducted to assess the significance of the detected GF. To this end, the likelihood was maximized under the constraint that *p*_*AC*_ and/or *p*_*BC*_ equalled zero. Twice the difference in log-likelihood between the unconstrained and constrained models was compared to a *χ*^2^ distribution with one (if a single parameter was set to zero) or two (if two parameters were set to zero) degrees of freedom. This procedure allows testing the significance of gene flow between *A* and *C*, of gene flow between *B* and *C*, or of gene flow generally.

### 2.6 Aphid: requirements, data filtering, running time

The Aphid program takes as input a set of rooted gene trees with branch lengths, a focal triplet of species and a set of outgroup species. This implies assuming that non-recombining segments including relevant phylogenetic information (*i*.*e*., several mutations) exist and have been correctly identified. Because branch lengths are usually expressed in per site number of changes, sequence length must also be input for each gene tree. Aphid only analyzes gene trees in which the three focal species are present and monophyletic and at least one outgroup species is present. When the considered gene tree contained several outgroup species, these were here required to be monophyletic. In addition, Aphid filters gene trees based on a criterion of clock-likeness. Specifically, the average distance from root to tips is calculated separately for the focal species, on one hand, and the outgroup, on the other hand, and the ratio between these two distances must be lower than a user-defined threshold, which was here set to 2. Aphid considers a gene tree as topologically unresolved (entailing specific likelihood calculation, see above) when the internal branch length is below a user-defined threshold, which was here set to 0.5 mutations. Gene trees in Aphid are supposed to describe the coalescence history of unlinked genomic segments, which implies assuming that no recombination has happened within a segment, and that recombination between segments is high. The segment size and between-segment distance satisfying these conditions are expected to vary among species depending on the effective population size and recombination landscape. Aphid typically takes less than a minute for analyzing data sets of thousands of gene trees on a laptop. This study used Aphid version 0.11.

## 3 Results

### 3.1 Simulations

Simulations were conducted in order to assess the performance of the Aphid method. First, data sets were simulated under the very Aphid model, parameters being set to *τ*_1_ = 0.01, *τ*_2_ = 0.015, *θ* = 0.005, *p*_*AB*_ = 0.2, *p*_*AC*_ = 0.1, *p*_*BC*_ = 0.05, *p*_*a*_ = 0.75. These defined the branch lengths and prior probability of each of the 14 possible scenarios (one no-event, three ILS, ten GF). To simulate a gene tree, a scenario was randomly picked, then branch lengths were drawn from Poisson distributions of means equal to the scenario expectations times sequence length times relative mutation rate, where sequence length followed a log-uniform distribution between 100 and 1000 bp, and relative mutation rate varied by a factor of nine among gene trees. A thousand data sets each made of 2000 gene trees were obtained this way and analyzed with Aphid. The results are shown in Supplementary Fig. 2. For all seven parameters, the method recovered the true value on average, as expected. The relative error rate (ratio of standard deviation to mean of the parameter estimate) was 0.03 for *τ*_1_, 0.02 for *τ*_2_, 0.06 for *θ*, 0.16 for *p*_*AB*_, 0.08 for *p*_*AC*_, 0.12 for *p*_*BC*_ and 0.05 for *p*_*a*_.

The performances of Aphid were furthered assessed under more challenging conditions, namely, the multi-species coalescent model with gene flow. I considered an ((*A,B*),*C*) ultrametric species tree with speciation time *t*_1_ between *A* and *B* and *t*_2_ between (*A, B*) and *C*, and population size *N*_*e*_. Migration was assumed to occur at constant rate *m*. Migration occurred between the three *A, B* and *C* species when time was lower than *t*_1_ (rate *m* for each of the three pairs of species), and between the ancestral (*A, B*) and *C* species when time was between *t*_1_ and *t*_2_, both ways. This implies that the simulations departed the Aphid scenarios in several respects. The space of simulated gene trees was continuous instead of a finite number of scenarios as assumed in Aphid, and covered a wider range of possibilities. For instance, very recent (*t < t*_1_*/*2) and ancient (*t > t*_1_) gene flow times were possible, as well as multiple events of gene flow.

A thousand data sets were simulated, each consisting of 2000 independently-generated gene trees. An outgroup distant from *A, B* or *C* by 2 million generations was added to every gene tree. Gene trees were assumed to correspond to genomic segments of length following a log-uniform distribution between 100 and 1000 bp. The mutation rate was assumed to vary by a factor of nine among genomic segments, with a mean value of 2 *×* 10^−8^ per generation per bp. The *t*_1_, *t*_2_, *N*_*e*_ and *m* parameters differed among the 1000 simulated data sets as they were drawn from uniform distributions: *t*_1_ took values in [0.5; 1.5] million generations, *t*_2_ − *t*_1_ in [0.25; 0.75] million generations, *N*_*e*_ in [50,000; 150,000] and *m* in [10^−8^, 10^−7^] per generation. The percentage of simulated gene trees with a discordant ((*A,C*),*B*) or ((*B,C*),*A*) topology) varied from 4% to 58% across data sets, with a mean of 28%.

The main results are presented in Fig.2, in which a random sample of 100 simulated data sets is shown for clarity. Fig.2A and B compare the estimated and simulated values of parameters *τ*_1_ and *τ*_2_ and population mutation rate *θ*. There was a tendency to slightly underestimate *τ*_1_ and *τ*_2_ and slightly overestimate *θ*. A similar bias was reported with the CoalHMM method (Dutheil et al., 2009), which also uses a discrete set of scenarios with fixed branch lengths. Fig.2C and D compare the estimated and simulated proportion of discordant gene trees resulting from ILS and GF, respectively. The prevalence of ILS (equation 6) and GF (equation 7) were reasonably well estimated under a wide range of conditions, particularly when the percentage of discordant gene trees was below 35% (closed dots, arbitrary threshold). For these low-conflict gene trees, Spearman’s correlation coefficient between true and predicted prevalence was 0.91 for ILS, 0.93 for GF (0.87 and 0.88, respectively, when all gene trees were considered). ILS tended to be slightly overestimated when its real effect was low, and underestimated when it was high.

**Figure 2.**
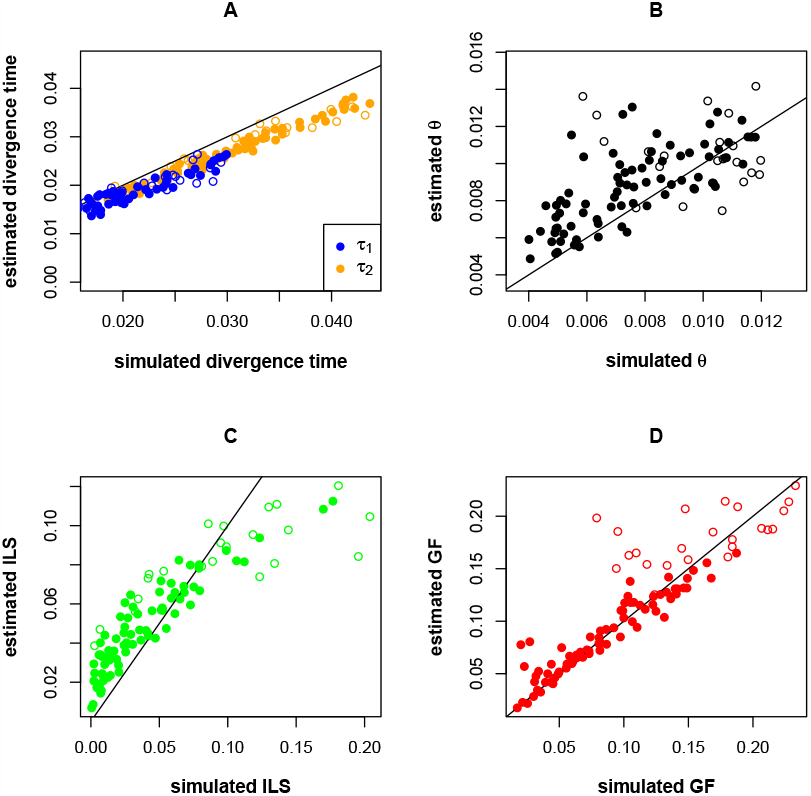
Simulation-based assessment of the performances of Aphid, multi-species coalescent model. A. Estimated *τ*_1_ and *τ*_2_. B. Estimated *θ*. C. Estimated ILS prevalence in phylogenetic conflicts. D. Estimated GF prevalence in phylogenetic conflict. True values are on the X-axis. Closed dots correspond to simulated data sets in which discordant topologies (*i*.*e*., simulated ILS + simulated GF) amounted 35% of the gene trees or less. A subset of 100 random data sets is shown, out of the 1000 simulated ones.

The QuIBL program (Edelman et al., 2019) was tested on these simulated data sets using the default options, with the exception of the numsteps parameter, which was set to 50 in agreement with the program documentation. QuIBL did not perform particularly well here, the estimated ILS and GF prevalence being only weakly correlated with the simulated ones (Supplementary Fig. 3A): Spearman’s correlation coefficient between true and predicted prevalence was 0.42 for ILS, 0.50 for GF. QuIBL assumes a constant mutation rate across loci, whereas the mutation varied substantially in our simulations. A second series of simulations with a constant mutation rate was therefore conducted. The correlation between the simulated and estimated GF/ILS contributions was improved, while not reaching the accuracy of Aphid (Supplementary Fig. 3B).

The performance of Aphid in annotating the status of a specific gene tree was also evaluated. For each locus of each simulated data set, the posterior probability of being affected by ILS (equation (4)) or by GF (equation (5)) was calculated, and the posterior probability of no-event was defined as one minus the sum of these two probabilities. If one of the three posterior probabilities was above 0.95 then the corresponding locus was annotated accordingly. If none was above 0.95 then no annotation was made. The percentage of annotated gene trees varied greatly among simulated data sets, from 4% to 76%, the average across data sets being 24%. Focusing on the annotation of simulated gene trees with a discordant ((*A, C*), *B*) or ((*B, C*), *A*) topology, it was found that 94% of the discordant gene trees predicted to result from GF actually experienced an event of GF in the simulations; only 6% of these were false GF positives, *i*.*e*., discordant gene trees having actually experienced ILS. The specificity was much lower as far as ILS-predicted discordant gene trees were concerned: only 54% of these were gene trees having truly experienced ILS in the simulations, the other 46% being false GF negatives. Results also accounting for ((*A, B*), *C*) gene trees are shown in Supplementary Fig. 4.

This result might be at least in part explained by the existence in the simulations of ancient events of GF, occurring close to *t*_1_ or earlier. The resulting gene trees have relatively long terminal branches, and are not properly taken into account by the Aphid GF scenarios. To test this hypothesis, an additional series of simulations was performed allowing GF to only occur before *t*_1_. Aphid performed similarly in terms of parameter estimation and GF discordant gene tree annotation. Among discordant gene trees predicted to be due to ILS, 82% indeed experienced ILS, and 18% were due to GF. This was a clear improvement compared to the simulations allowing for ancient GF, confirming that Aphid probably struggles distinguishing ILS from ancient GF. Of note, the bias in parameter estimation visible in Fig.2A and B was still observed in these simulations lacking ancient GF.

The above simulations were conducted under a model in which GF affects the A and B lineages to the same extent. An additional series of simulations was performed, in which GF was asymmetric across lineages. Specifically, GF between B and C was assumed to occur with a probability ten times as high as GF between A and C. In terms of parameter estimation and ILS/GF detection Aphid had performances similar to the symmetric scheme. The estimated index of imbalance associated with ILS or GF (equation 8 and 11) was compared among the two schemes. Under the symmetric GF scheme, the estimated imbalance was close to 0.5 for the two processes (mean 0.52 and 0.54 for ILS and GF, respectively). Under the asymmetric scheme, the mean estimated imbalance was 0.53 as far as ILS was concerned, and 0.83 as far as GF was concerned, confirming that under these conditions Aphid is able to correctly distinguish between the two processes and detect GF asymmetry.

Overall these simulations suggest that (i) Aphid estimates the relative contribution of ILS and GF to phylogenetic conflicts with reasonably good accuracy across a wide range of conditions, provided the overall conflict prevalence is not too high, and (ii) Aphid only annotates a limited fraction of genes (with the sequence length and mutation rate considered here), while calling a small number of false GF positives.

### 3.2 Exon tree analysis

The Aphid program was applied to three primate data sets downloaded from the OrthoMaM v10 database (Scornavacca et al., 2019). These data sets, depicted in Supplementary Table 2, correspond to triplets of species from genus Macaca (macaques), tribe Papionini (baboons and allied), and subfamily Homininae (African apes). Speciation times for species in these triplet range from 3.3 to 8.6 Mya according to the TimeTree of Life (http://www.timetree.org/). Gene trees reconstructed from alignments of exons of length 400 bp or above were selected — exons were previously identified as appropriate gene tree reconstruction units in mammals (Scornavacca and Galtier, 2017). There were 2687 (Macaca), 2677 (Papionini) and 3398 (Homininae) eligible exon trees per data set after filtering for taxon sampling and clock-likeness (see above 2.6). In all three data sets a majority of the gene trees agreed with the OrthoMaM reference species tree topology (https://orthomam.mbb.cnrs.fr/). The percentage, among resolved gene trees, of discordant topologies were 20% (Macaca), 38% (Papionini) and 25% (Homininae), and the average sequence divergence varied from 0.21% to 0.92%.

Dot size in Fig.3 reflects the estimated contributions of ILS (green, equation 8) or GF (red, equation 11) to the conflict, which are also given explicitly as numbers. The two processes contributed a substantial fraction of the conflict in the three data sets. In all three cases a model assuming no GF (parameters *p*_*AC*_ and *p*_*BC*_ fixed to zero) led to a significant drop in log-likelihood and was rejected with a *p*-value below 10^−10^ by a likelihood ratio test. The Y-axis in Fig.3 is the estimated imbalance between the two discordant topologies associated with the detected effects. As far as ILS was concerned, the imbalance was low in the three data sets, as expected. The estimated GF was associated with strong discordant topology imbalance in Macaca. In this data set, Aphid detected significant GF between the two Indonesian species *Macaca fascicularis* and *Macaca nemestrina*, but much less GF between *Macaca mulatta* and *M. nemestrina*, consistent with published ABBA-BABA-based analyses (Vanderpool et al., 2020; Song et al., 2021). This in essence says that in Macaca, the ((*fascicularis, nemestrina*), *mulatta*) and ((*mulatta, nemestrina*), *fascicularis*) topologies appear in roughly equal numbers as far as tall discordant gene trees are concerned, whereas the former topology is highly dominant among short discordant gene trees. Note that the signal captured by ABBA-BABA is averaged across all the discordant gene trees, irrespective of branch lengths. Black squares in Fig.3 show the magnitude of this merged imbalance, which in Macaca was much weaker than the GF-specific imbalance. By separating the signal from tall *vs*. short gene trees, Aphid appears to provide a more informed description of the situation.

**Figure 3.**
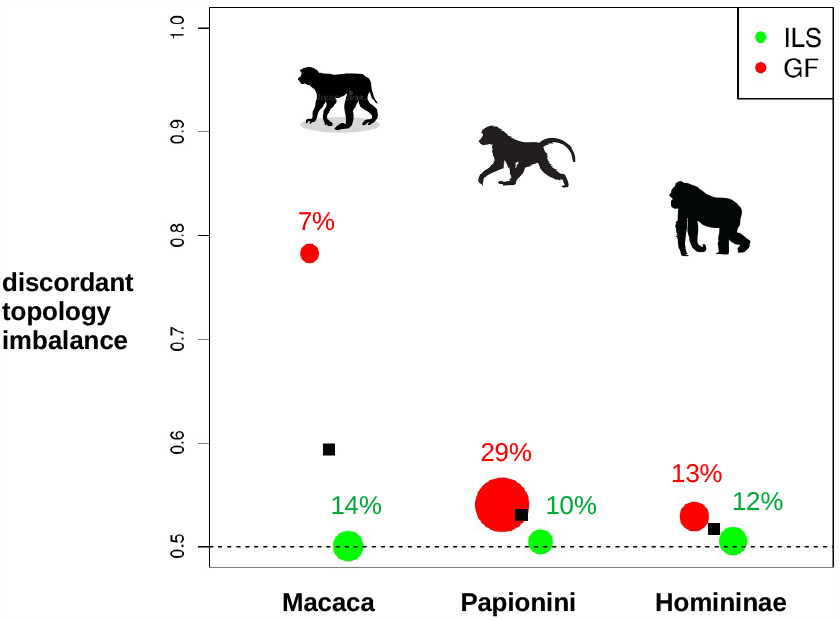
Aphid analysis of primate exon tree data sets. Dot size reflects the estimated contribution of ILS (green) or GF (red) to the phylogenetic conflict, the actual estimate being given explicitly near each dot. Y-axis: imbalance between the contributions of the ((A,C),B) and ((B,C),A) topologies to the detected effect. Black squares: overall imbalance, merging the ILS and GF signals.

Interestingly, in the other two data sets, Aphid detected substantial gene flow despite low discordant topology imbalance — a signal necessarily missed by ABBA-BABA. This was notably the case in the Homininae data set, *i*.*e*., the ((human,chimpanzee),gorilla) triplet. This triplet has been thoroughly studied during the last decades, with the conflict most often interpreted in terms of ILS (Hobolth et al., 2007; Dutheil et al., 2009; Rivas-González et al., 2023a). Aphid suggests instead that roughly half of the discordant exon trees are shorter than expected under ILS, and more likely to result from chimpanzee ↔ gorilla and human ↔ gorilla gene flow. This was a surprising result that I aimed at confirming via the analysis of non-coding data.

### 3.3 Non coding alignment analysis

Mammalian non-coding sequence alignments were downloaded from the UCSC server https://genome.ucsc.edu/. These alignments have been obtained based on the genome sequence of 30 species, among which only the 20 anthropoid primates were considered. Alignments of length >500 bp comprising all of the following eight species were selected: *Homo sapiens, Pan troglodytes, Gorilla gorilla, Pongo abeli, M. mulatta, Papio anubis, Callithrix jacchus, Saimiri boliviensis*. For each alignment, a phylogenetic tree was built using default parameters in IQTree. Trees were rooted in a way that placed the (*C. jacchus, S. boliviensis*) clade — i.e., Platyrrhini — as an outgroup to the rest of the species. When *C. jacchus* and *S. boliviensis* did not form a phylogenetic group, trees were instead discarded. This resulted in 495,614 gene trees corresponding to alignments of average length 761 bp, of which 95% had a length between 508 and 1584 bp.

Aphid was run using ((human,chimpanzee), gorilla) as the focal triplet and Cercopithecidae (*M. mulatta, M. fascicularis, P. anubis, Chlorocebus sabeus, Rhinopithecus roxellana, Nasalis larvatus*) as the outgroup, the other species being disregarded. The average sequence divergence was 1.1% between human and chimpanzee, 1.5% between human and gorilla. The percentage of discordant topologies was 29%, of which 49% grouped human with gorilla and 51% grouped chimpanzee with gorilla. We first analyzed chromosomes separately, meaning 23 Aphid runs — 22 autosomes and the X chromosome. Autosome data sets included an average 22,528 gene trees (from 3,779 in chromosome 19 to 49,164 in chromosome 2), while the X chromosome data set included 22,197 gene trees. The results are provided in Supplementary Table 3 and summarized in Fig.4, in which the top six panels show the distribution of parameter estimates across chromosomes, and the two bottom panels the distribution of the estimated ILS and GF prevalence. Estimates of parameter *p*_*a*_ (not shown in Fig.4) varied between 0.94 and 0.99 among chromosomes.

**Figure 4.**
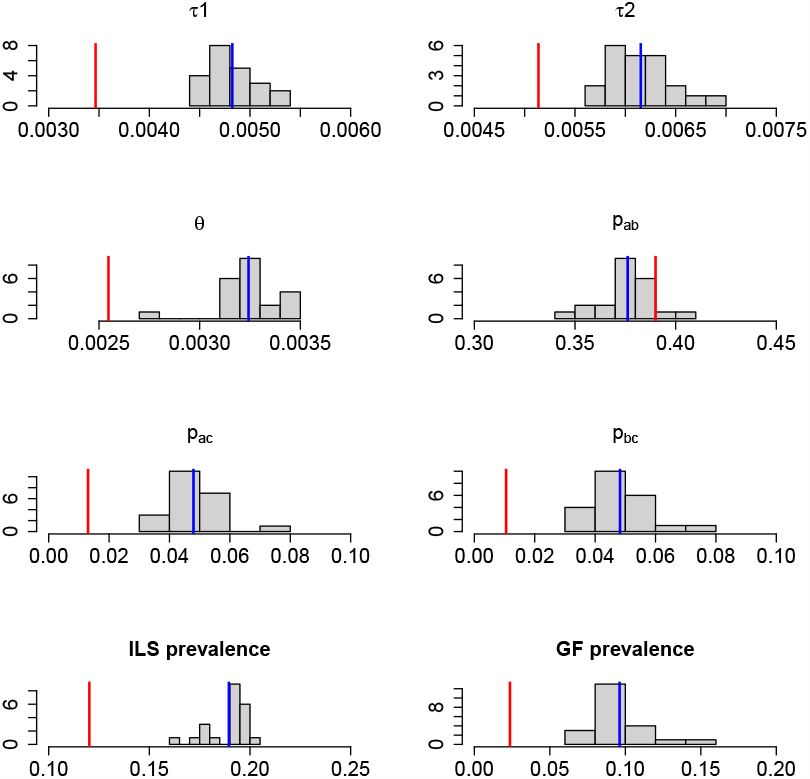
Aphid analysis of non-coding alignments in Homininae. The top six histograms show the distribution of parameter estimates across autosomes. The bottom two histograms show the distribution of the estimated contribution of ILS, respectively GF, to the phylogenetic conflict across autosomes. Blue vertical bar: average among autosomes. Red vertical bar: X chromosome estimate.

In autosomes, the mean estimated prevalence of phylogenetic discordance due to ILS was close to 18%, which was higher than the 12% obtained based on exon trees (see above Fig.3). This difference was expected since coding sequences are presumably more strongly affected than non-coding regions by linked selection, which tends to reduce coalescence times and consequently decrease the probability of ILS (McVicker et al., 2009; Hobolth et al., 2011). The estimated contribution of GF to the phylogenetic conflict was 9.6%, which is a bit lower than estimated from coding sequences, but still substantial — and highly significant for all chromosomes. Non-coding sequence analysis therefore corroborated the results obtained from exon trees, based on a much larger data set. The mean estimated *p*_*a*_ parameter equalled 0.95 for the Homininae data set, meaning that most of the detected gene flow is inferred to have happened shortly after the human|chimpanzee split. Please keep in mind, however, that we are here talking about the time of coalescence of lineages brought together by GF, which is expected to be older than the time at which GF occurred.

The X chromosome (red vertical bars) was a clear outlier. The prevalence of both ILS and GF were estimated to be weaker, and the estimated *τ*_1_, *τ*_2_ and *θ* lower, on the X than on autosomes. These results are consistent with the hypotheses that, compared to autosomes, the X chromosome experiences a lower mutation rate, a lower *N*_*e*_, and is more resistant to introgression, as expected theoretically and previously reported in a number of studies (Hobolth et al., 2007; Makova and Li, 2002; Wilson and Makova, 2011; Hammer et al., 2008; Keinan et al., 2009; Ellegren, 2009; Amster et al., 2020; Geraldes et al., 2008; Sankararaman et al., 2014).

Two hundred random samples of 10,000 autosomal gene trees were generated and analyzed with Aphid, thus providing a direct, empirical measurement of the sampling variance associated with parameter estimation. The first five rows of Supplementary Table 4 provides the estimated parameters and confidence intervals for five replicated data sets out of 200. The sixth row of Supplementary Table 4 provides the mean parameter estimates across replicates, as well as 95% intervals obtained by excluding the 2.5% smallest and 2.5% largest values. Comparison of the sixth row with any of the first five suggests that the confidence intervals returned by Aphid correctly capture the sampling variance associated with parameter estimation. These confidence intervals, however, are typically narrower than suggested by the chromosome-by-chromosome analysis (Fig.4). The seventh row of Supplementary Table 4 indicates, for each parameter, the number of autosomes with an estimate outside the 95% interval calculated based on replicates. There are 22 autosomes, so the expectation here would be 1.1, but the observed numbers were always higher. This suggests that the gene trees carried by a particular autosome tend to have correlated branch lengths. At any rate, it should be recalled that the sampling variance is not the sole source of uncertainty in Aphid-based inferences, departure from the model assumptions being another important issue to consider (see discussion).

Using whole-genome data and empirical estimates of the mutation rate and generation time, Rivas-González et al. (2023a) dated the average coalescence time between human and chimpanzee at 10Mya (their Fig. 1A). The average sequence divergence between human and chimpanzee in our data set is 1.15 *×* 10^−2^ per bp. Combining these two numbers, one gets an estimated neutral mutation rate of 5.75 *×* 10^−4^ per bp per My for our data. Dividing the estimated *τ*_1_ and *τ*_2_ by the mutation rate yields an estimated human|chimpanzee speciation time of 8.3My (95% confidence interval: [8.1; 8.5]) and an estimated human|gorilla speciation time of 10.7My (95% confidence interval: [10.5; 10.8]). Assuming a generation time of 20 years for apes, one gets a per bp per generation neutral mutation rate of 1.15 *×* 10^−8^. Dividing our estimated *θ* by four times this figure yields an estimated effective population size of 70,300. Rather using a human-gorilla coalescence calibration, dated at 12.4 Mya by (Rivas-González et al., 2023a), provides younger speciation time estimates (human|chimpanzee: 7.6 Mya; human|gorilla: 9.8 Mya) and a smaller estimated effective populatiopn size of 64,500. Assuming that the phylogenetic conflict entirely results from ILS, Rivas-González et al. (2023a) rather reported estimated speciation times of 6 Mya (human|chimpanzee) and 8 Mya (human|gorilla), and ancestral effective population sizes of 177,000 (human-chimpanzee ancestor) and 107,000 (human-gorilla ancestor).

## 4 Discussion

Aphid models a set of gene trees as a mixture of typical scenarios, and returns an estimate of the contribution of the various (categories of) scenarios to the data. The analysis of simulated and real data suggested that Aphid is reasonably robust to a wide range of conditions and has the potential of revealing gene flow despite balanced discordant topology counts, thus extracting more information from the data than the simpler ABBA-BABA.

### 4.1 Aphid: strengths and limitations

Likelihood calculation in Aphid is approximate in that it does not account for the variance in coalescence time among loci. The model assumes that gene trees associated with a given scenario only differ from each other due to the stochasticity of mutation, neglecting the variance due to drift. There are two connected reasons why this approximation is made. First, the distribution of branch lengths under the multi-species coalescent is uneasy to write down and integrate (Takahata et al., 1995; Yang, 2010). Secondly, the calculation and maximization of the approximated likelihood are achieved efficiently, since the likelihood can be analytically differentiated with respect to parameters. The Aphid method typically takes a few seconds to analyse a data set of several thousand gene trees, offering possibilities such as data re-sampling for instance. Our simulations suggest that inferences are reasonably robust despite the approximation. Yet, one should keep in mind that Aphid is a heuristic and does not rely on a proper population genetic model.

The set of scenarios used by Aphid is a simplified representation of the actual complexity of gene trees. In particular, Aphid assumes that two lineages brought together by GF coalesce in the donor species earlier than the next speciation event (thinking backwards in time). In reality, this coalescence could be older than *t*_1_ or *t*_2_, which entails complex scenarios in which GF and ILS interact. Aphid is based on the premise that GF and ILS leave distinguishable signatures on gene tree branch lengths, which is only valid if GF is sufficiently recent. Phylogenetic conflict due to ancient GF — *i*.*e*., GF time of the order of speciation time — is unlikely to be distinguished from phylogenetic conflict due to ILS by Aphid, or by any method analysing samples of one lineage per species. If the species history is such that scenarios in which ILS and GF co-occur are numerous, this is expected to fault Aphid, which in essence is based on the assumption that the two processes are disjoint. One way of mitigating this problem is to refrain using Aphid, or be careful with the interpretation, when the overall level of conflict is too high — say, above 50 or 55% of discordant topologies.

As Aphid aims to interpret branch lengths in terms of coalescence times, properly modeling the among loci variation in mutation rate is essential. The strategy retained here consists in (1) including an outgroup, (2) removing gene trees departing from the molecular clock hypothesis, (3) estimating a locus relative mutation rate as the average root-to-tip distance. Although plausible this procedure could probably be improved. Step (2) above, in particular, implied discarding up to 40% of exon trees in our analysis of primate data. Instead, one might explicitly model the among lineage rate variation and estimate the triplet mutation rate even for gene trees departing from the molecular clock assumption (Thorne et al., 1998; Lartillot et al., 2009). Another potential issue with the above procedure is that the average root-to-tip distance depends not only on mutation rate but also on the age of the root node, which is expected to differ among loci due to the variance of the coalescent, and possibly ancient GF. For this reason I recommend using a relatively distant outgroup, such that the standard deviation in root age is small compared to root age. It should be noted that the uncertainty in locus-specific mutation rate estimation was not taken into account in the confidence intervals calculated here.

Aphid in essence intends to distinguish short from tall discordant gene trees, annotating the former as GF and the latter as ILS. This, however, might be a simplistic interpretation of the variation of gene tree height among loci. Old coalescence times, in particular, are not necessarily explained by ILS, but could reflect GF from extinct or unsampled anciently diverged lineages (Green et al., 2010; Pease and Hahn, 2015; Tricou et al., 2022a). A recent simulation study suggests that GF from so-called “ghost” lineages could be pervasive and underestimated (Tricou et al., 2022b). If GF from ghost is asymmetric, this might manifest itself in Aphid by an *I*_*ILS*_ index significantly different from 0.5 (equation 8 and 11). If however GF from ghost affects the A and B lineages to the same extent, then the signal will probably be indistinguishable from ILS in Aphid.

Aphid makes a number of approximations, among which the expected branch lengths under the no-event scenario (Fig. 1). Here the first coalescence is assumed to happen at time *t*_1_ + 2*N*_*e*_, which is the mean coalescence time under the assumption of no GF. The no-event scenario, however, assumed no GF and no ILS, meaning that the first coalescence time should logically be constrained to be more recent than *t*_2_, implying a mean lower than *t*_1_ + 2*N*_*e*_. The approximation is valid if 2*N*_*e*_ ≪ *t*_2_ − *t*_1_, *i*.*e*., if ILS is negligible, which is not generally true. It is interesting to note that Aphid performed reasonably well in the simulations conducted here even though *N*_*e*_ was not generally much smaller than *t*_2_ − *t*_1_.

### 4.2 Relationships with other methods

Aphid has a lot in common with the CoalHMM method (Hobolth et al., 2007; Dutheil et al., 2009), which also uses discrete categories of scenarios with branch lengths equal to theoretical expectations. CoalHMM does not analyse supposedly independent gene trees but models the topological variation across a genome — a more ambitious objective. By explicitly modeling recombination, CoalHMM uniquely offers the opportunity of comprehensively analyzing genome-wide data without the need of identifying unlinked, non-recombining genomic windows (see below). Another major difference is that CoalHMM does not consider GF, and rather assumes that the topological conflict entirely results from ILS — not an insignificant assumption. The ABBA-BABA-based literature has reported a significant imbalance between discordant topologies in many taxa, suggesting that GF is not a generally negligible source of phylogenetic conflict. Interestingly, the same bias in parameter estimation found with Aphid (underestimated speciation times, overestimated *θ*) was reported with CoalHMM (Dutheil et al., 2009), and this problem was recently solved by including additional ILS scenarios covering a wider range of branch lengths (Rivas-González et al., 2023b) — an improvement that might be worth implementing in Aphid as well.

Aphid also has similarities with the maximum-likelihood method proposed by Yang (2010). Like Aphid, this method analyses supposedly independent three-species gene trees and models the among loci variation in mutation rate. Unlike Aphid and CoalHMM, it numerically integrates across all possible coalescence times, thus providing an exact likelihood calculation, while also accounting for multiple mutations at the same site. A major difference is that the method introduced by Yang (2010) models GF between the two most recently diverged species — *A* and *B* according to our notations — whereas Aphid mainly focuses on GF between *A* and *C* or *B* and *C*. Detecting GF between *A* and *B* is a difficult problem since this does not entail any topological change. Yang (2010) proposes a test of the null hypothesis of an absence of GF, drawing information from the extra variance in *A*|*B* coalescence time expected in case of GF. A similar analysis is in principle possible in Aphid by testing the significance of the *p*_*AB*_ parameter. Unlike Yang’s method Aphid was not specifically designed for this goal, and its performance with this respect was not assessed in the current study.

Like Aphid, Legofit (Rogers, 2019) is a maximum-likelihood method explicitly modeling both GF and ILS. Legofit is more general and ambitious than Aphid, allowing an arbitrary number of species/populations and a diversity of models of population history. Unlike Aphid, however, Legofit only considers one-directional gene flow, in the spirit of ABBA-BABA approaches. Because it takes information from site patterns, Legofit does not need to make any assumption regarding recombination (see below). In its initial version Legofit was computationally demanding, but recent developments have considerably improved its efficiency (Rogers, 2022). So far Legofit has mostly been used for characterizing archaic introgression among human lineages, or among chimpanzee lineages (Brand et al., 2022). It would be great to investigate whether it can be applied to more ancient divergences.

The published method maybe most similar in spirit to Aphid is QuIBL (Edelman et al., 2019). Like Aphid, QuIBL aims at partitioning the discordant gene trees into GF and non-GF categories via a mixture model. Unlike Aphid, QuIBL is an exact method in that it properly accounts for the among loci variance in coalescence time. This, however, comes at the cost of only focusing on a single branch, namely the internal branch (*d* according to our notations), the distribution of which can be analytically predicted conditional on the topology. QuIBL did not perform particularly well in the simulations conducted here. This is in part explained by the fact that the mutation rate varied substantially across loci in our simulations, whereas QuIBL assumes a constant mutation rate (Supplementary Fig. 3). It should also be noted that relatively short sequences were simulated — 100-1000 bp, to be compared to windows of 5 kb in Edelman et al. (2019). Because it analyzes only part of the available information, QuIBL likely requires more data than Aphid to achieve the same accuracy. One, however, should not conclude from this study that Aphid is generally superior to QuIBL. Further work is needed to characterize the applicability conditions for both methods.

### 4.3 The no intra-locus recombination assumption

Like all gene tree-based methods Aphid implicitly assumes that each of the analyzed gene trees was built from a piece of DNA unaffected by recombination. It is important to ask whether this assumption is valid, and if not, to what extent it is a problem. I here focus on the Homininae situation. In humans, the per bp, per generation average recombination rate was estimated to be 1.25 *×* 10^−8^ (Jensen-Seaman et al., 2004). This implies that a segment of length 800bp (roughly the average alignment length in our non-coding data set) is expected to be transmitted as a single block across 10^5^ generations, on average. The human-gorilla speciation dates back to ∼10My, which is 5 *×* 10^5^ generations assuming a generation time of 20 years. So clearly the segments we are looking at have not traversed the Homininae phylogeny as blocks, but have been affected by an average ∼10-15 events of recombination.

Not every recombination event, however, is expected to affect the observable variation, and especially so when one considers isolated populations/species (Lanier and Knowles, 2012). Thinking backwards in time, recent recombination events have a small probability of being effective since the recombining lineages are likely to coalesce with one another before reaching speciation time — see figure 4 in Xu and Yang (2016). The simulations conducted by Zhu et al. (2022) revealed a surprisingly small effect of intra-locus recombination on inferences under the multi-species coalescent model, gene trees from neighboring segments being strongly correlated. Although the existing literature on the subject is mostly empirical, it appears reasonable to postulate that the proportion of recombination events effectively affecting gene trees in the human/chimpanzee/gorilla case is well below 0.5 — *e*.*g*. see Zhu et al. (2022). Another important element to consider is the existence of recombination hot spots. In humans, it was estimated that roughly 80% of the recombination is concentrated in 15-20% of the genome (Myers et al., 2005). This implies that 80-85% of the loci from which gene trees were built have experienced a recombination rate roughly four times as small as the genome average. For these loci, and given the above, the assumption of an absence of intra-locus recombination does not seem to be of much concern.

The high-recombining fraction of our data sets, however, probably deviates from the no-recombination assumption, meaning that, for these loci, what we call a gene tree is actually some kind of consensus across distinct genealogies. The consequences on the behaviour of Aphid are not easy to predict. I suspect that overlooking intra-locus recombination should tend to reduce the observed phylogenetic conflict, and thus mainly limit our power to detect GF or ILS. Yet one cannot exclude that intra-locus recombination might distort the estimated branch lengths in a way that biases the inference (Schierup and Hein, 2000). Additional simulations might help here. Finally, it should be noted that the above rationale implies that, at least in primates, the appropriate segment length to be used for gene tree reconstruction is of the order of 100-1000bp. This implies that any method aiming at analyzing such data should be ready to face non-negligible sampling error in topology and branch length measurement — and see above discussion of the Aphid/QuIBL comparison.

### 4.4 Ancient gene flow in apes

Application of Aphid to coding and non-coding data in great apes provided strong support for the existence of ancient GF between human, chimpanzee and gorilla shortly after the human/chimpanzee split. This is a novel result, based on the discovery that a substantial fraction of discordant gene trees have shorter, not longer, terminal branches, compared to canonical ((human, chimpanzee), gorilla) genealogies. Analyzing the genome-wide distribution of the human/chimpanzee sequence divergence, Patterson et al. (2006) suggested that the speciation between these two species was complex and involved some GF, but this claim was criticized (Wakeley, 2008; Presgraves and Yi, 2009). Subsequent analyses sometimes did (Yang, 2010) and Mailund et al. (2012) and sometimes did not (Yamamichi et al., 2012) reject the null hypothesis of an absence of GF between human and chimpanzee. Indeed, inferring GF from the analysis of just one sequence in each of two species is a difficult task given the many forces governing the distribution of pairwise sequence divergence — mutation, demography, selection. When more than two species were used, a model assuming no GF was most often applied in great apes, presumably because ABBA-BABA-like approaches did not reveal any asymmetry among discordant topologies. The existence of ancient GF in this group, however, comes as no surprise. GF is a pervasive process that accompanies genome divergence during speciation and is more or less systematically detected when sequence divergence is low enough (Roux et al., 2016). There is no obvious reason why great apes should be an exception.

Besides the interpretation of the phylogenetic conflict, explicitly modeling GF is likely to also make a difference in estimating parameters, particularly speciation times and ancestral effective population sizes. To assume no GF when there is some is expected to result in underestimated speciation times — due to recent GF-induced coalescence — and overestimated ancestral *N*_*e*_ — due to GF-induced discordance (Leaché et al., 2014). Indeed, in apes the speciation times estimated by Aphid are older, and the ancestral *N*_*e*_ smaller, than estimates obtained by the CoalHMM method (Rivas-González et al., 2023a), which only considers ILS as a potential source of conflict. The Aphid speciation time estimates are consistent with a recent review of the palaeontological data in apes (Almécija et al., 2021), in which the human|chimpanzee split was dated at 7.5 Mya and the human|gorilla split at 11 Mya. The Aphid ancestral *N*_*e*_ estimate is close to current estimated *N*_*e*_ in humans, chimpanzee, bonobo and gorilla (20,000-60,000 according to Supplementary Table S2 in Rivas-González et al. (2023a)).

### 4.5 Potential improvements

Aphid makes a number of assumptions that might limit its potential. GF in Aphid is assumed to be bidirectional, with an equal probability of GF in either way. The method therefore does not account for the kind of scenarios most usually considered when discussing GF between *H. neanderthalensis* and *H. sapiens*, for instance. Like QuIBL, Aphid assumes a constant *N*_*e*_ throughout the whole species tree, whereas CoalHMM and Yang’s method allow the (*AB*) ancestral *N*_*e*_ to differ from the (*ABC*) ancestral *N*_*e*_. Simulations indicate that the CoalHMM method is able to correctly estimate the two distinct ancestral *N*_*e*_’s when there is no GF (Dutheil et al., 2009). Intuitively, it seems plausible that, in the absence of GF, the minimal sequence divergence between two genomes informs on the speciation time, and its variance on the ancestral *N*_*e*_. In models accounting for GF, such as Aphid and QuIBL, the variance in coalescence time does not only depend on the ancestral *N*_*e*_ but also on the time and amount of GF, perhaps raising identifiability issues (Zhu and Degnan, 2017; Yang and Flouri, 2022). To assume a constant *N*_*e*_ when GF is modelled, even if biologically less realistic, might therefore appear safer — a formal argument on this subject would be welcome, though.

## 5 Conclusion

Aphid is an efficient three-species method quantifying the contribution of gene flow and incomplete lineage sorting to the phylogenetic conflict, while providing gene flow-aware estimates of speciation times and ancestral effective population size.

## 6. Acknowledgments

The author thanks Iago Bonicci and Khalid Belkhir from the Montpellier Bioinformatics & Biodiversity platform, Richard Durbin, Julien Dutheil, Julien Joseph, Alan Rogers, Marjolaine Rousselle, Céline Scornavacca, Carole Smadja, Ziheng Yang and two anonymous reviewers for comments, advice and help.

## 7 Supplementary Material, Data accessibility

Aphid and documentation are available from https://gitlab.mbb.cnrs.fr/ibonnici/aphid. The supplementary figures, supplementary tables, data sets and scripts used in this study are available from https://doi.org/10.48579/PRO/F6KEDM.

