## Supplementary tables and figures for "An approximate likelihood method reveals ancient gene flow between human, chimpanzee and gorilla": Supp_Fig4.pdf (from another version of this preprint: version 2, 2023-11-22)

### A. All gene trees

simulated (%) / predicted (%)

| predicted (%) | NoE | ILS | GF |
|---|---|---|---|
| NoE | 93.5 | 0.4 | 6.2 |
| ILS | 76.0 | 12.1 | 11.9 |
| GF | 21.3 | 4.3 | 74.4 |

### B. Discordant gene trees

simulated (%) / predicted (%)

| predicted (%) | ILS | GF |
|---|---|---|
| ILS | 54.5 | 45.5 |
| GF | 5.8 | 94.2 |
