## Supplementary figures and images for "An approximate likelihood method reveals ancient gene flow between human, chimpanzee and gorilla" (from another version of this preprint: version 2, 2023-11-22)

### Supp_Fig1.pdf

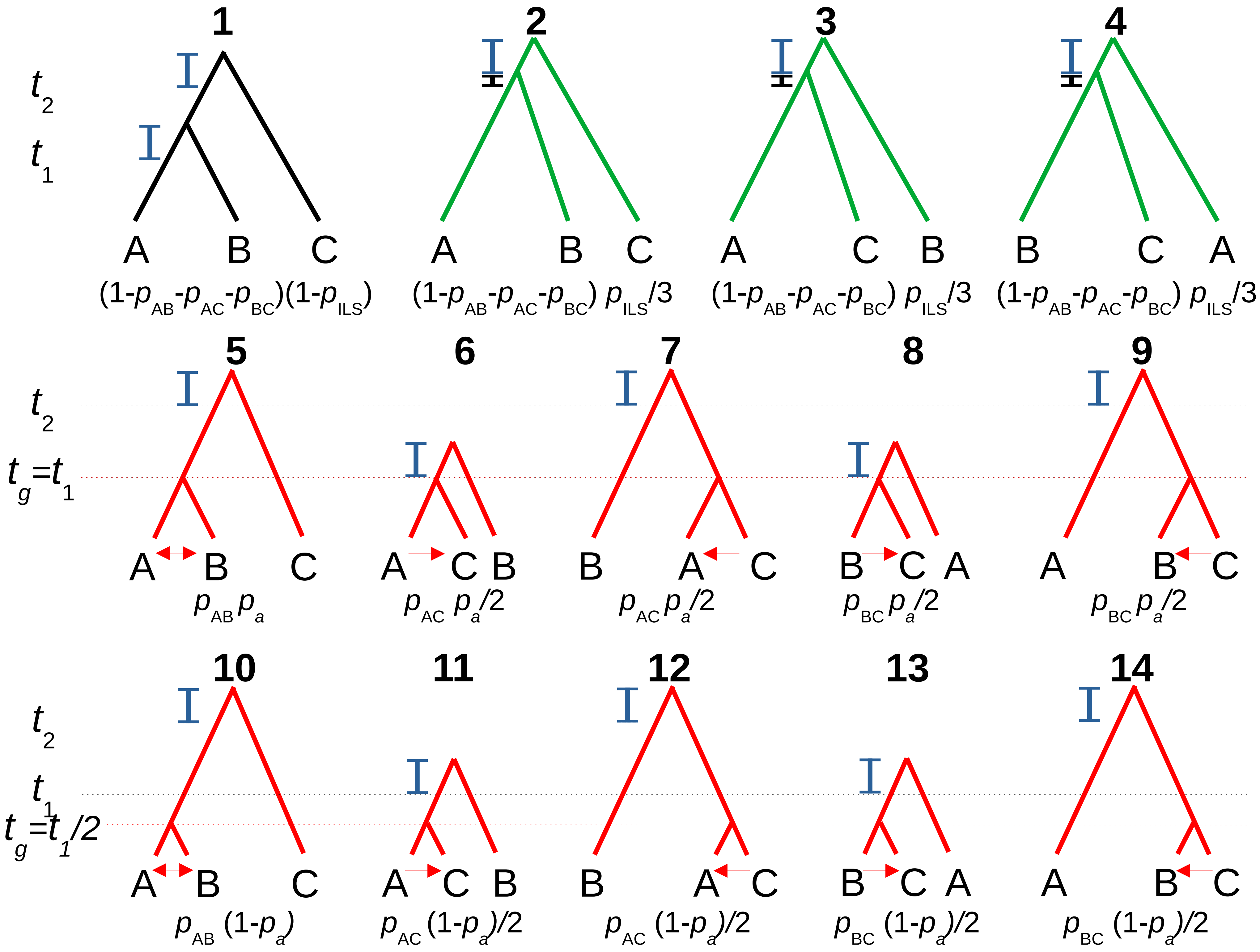

### Supp_Fig2.pdf

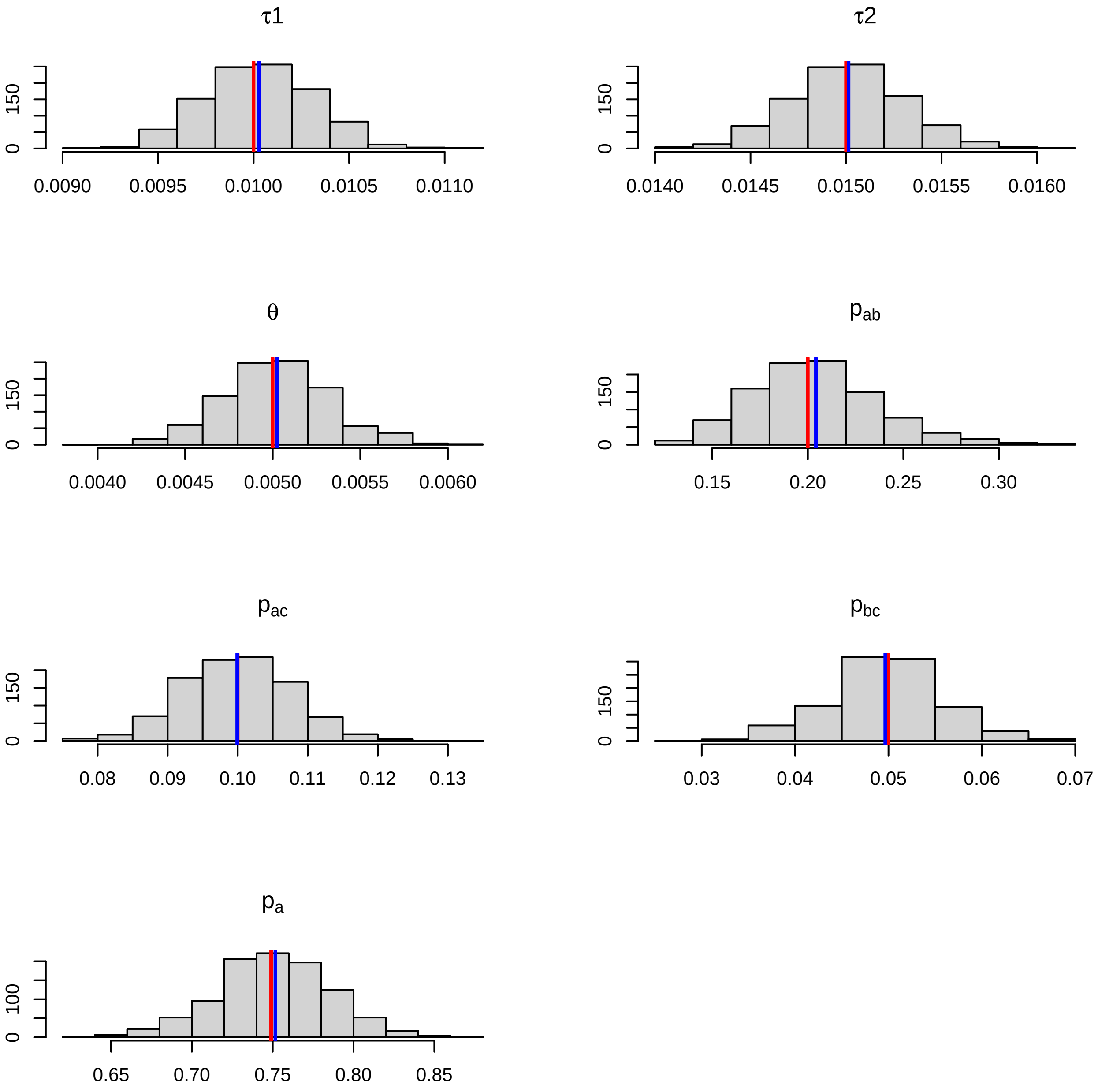

### Supp_Fig3.pdf

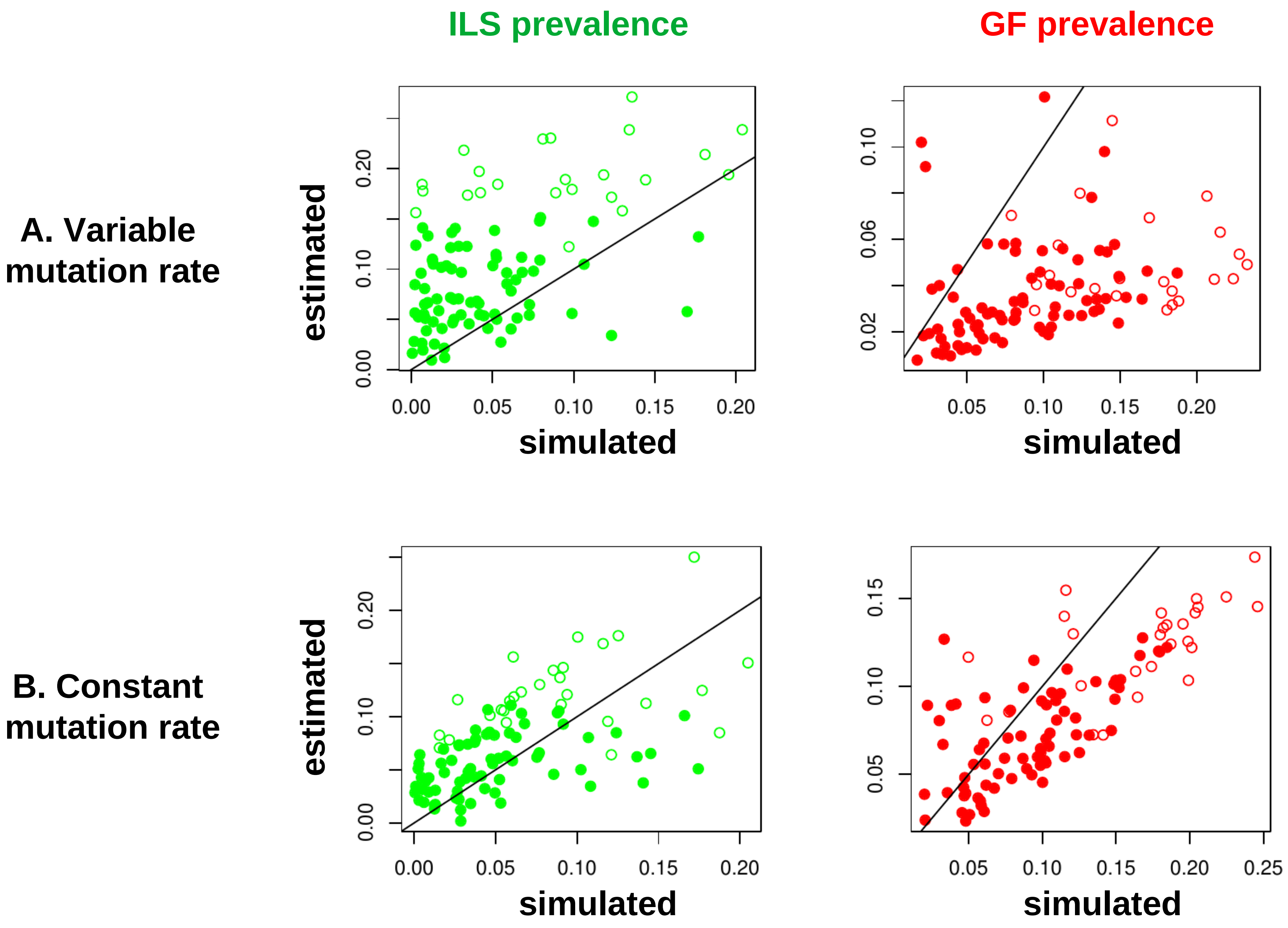
